# Accelerated tissue repair through cell proliferative effect by a size controlled aqueous based Fullerene C_60_ nanoformulation

**DOI:** 10.1101/2020.07.03.181537

**Authors:** Nabodita Sinha, Avinash Y. Gahane, Ashwani Kumar Thakur

## Abstract

We have developed Fullerene-C_60_ nanoformulations containing discrete sized nanoparticles by dispersing concentration range of Fullerene. Small sized particles are cytotoxic while larger ones are cell proliferative. The cell proliferative property is used for tissue repair in cellular and animal wound models.

---

The complex interplay between cells and nanoparticles changes with nanoparticle type, dispersion methods, and cell/tissue type. (1) Several nanoparticles are projected for anticancer, antimicrobial, bioimaging and smart polymer applications. (2,3) Fullerene (C_60_) allotrope has shown promise in cosmetic usage, dietary supplements and as an antitumor agent. These properties are primarily linked to its high antioxidant response and photodynamic cytotoxic effect on tumors (4–7). Nevertheless, potential of Fullerene for nanomedicine development is not yet fully explored, partly due to dispersion problems of pristine Fullerene. Attempts in the past were made to disperse C_60_ in tetrahydrofuran (THF). This dispersion produced cell death due to THF-intermediate formation as a by-product. (8) When dispersed in Tween 80, Fullerene showed no apparent toxicity. In olive oil, it increased the lifespan of rats. (9,10) On the other hand, in corn oil and saline water it showed genotoxicity. While corn oil has some inherent toxicity, but similar level of toxicity in saline water suggested that, Fullerene may also show cytotoxic effects. (11) Due to ease of dispersing in aqueous media, water soluble derivatives of Fullerene are also explored. Owing to differential derivatization properties these produced inconsistent reports for biological activities in some cases. (12) Some attempts were also made to disperse Fullerene C_60_ in water and cell culture media. Several groups also used solubilization of Fullerene in toluene and exchanging it with water. (13) However, detailed physicochemical characterization of the nanoparticles and effective concentration usage was not studied to understand the biological effects in some dispersion systems. (14) In this communication, we show that pristine Fullerene in different concentration ranges can be dispersed in physiologically relevant aqueous media. The resulting composition is denoted as nanoformulation.

We attempted dispersion of Fullerene C_60_ powder in water, phosphate buffer saline (PBS), DMEM, and DMEM+10% FBS by stirring at 700 rpm for 14 days at 25°C. To test the dispersion systems, 0.1, 10, 100, and 1000 μg/mL concentrations were chosen. Except for DMEM+FBS (**Figure 1 a**), all the other mediums (Water, PBS, DMEM) showed a high polydispersity index and micrometer-scale particle size.

**Figure 1:**
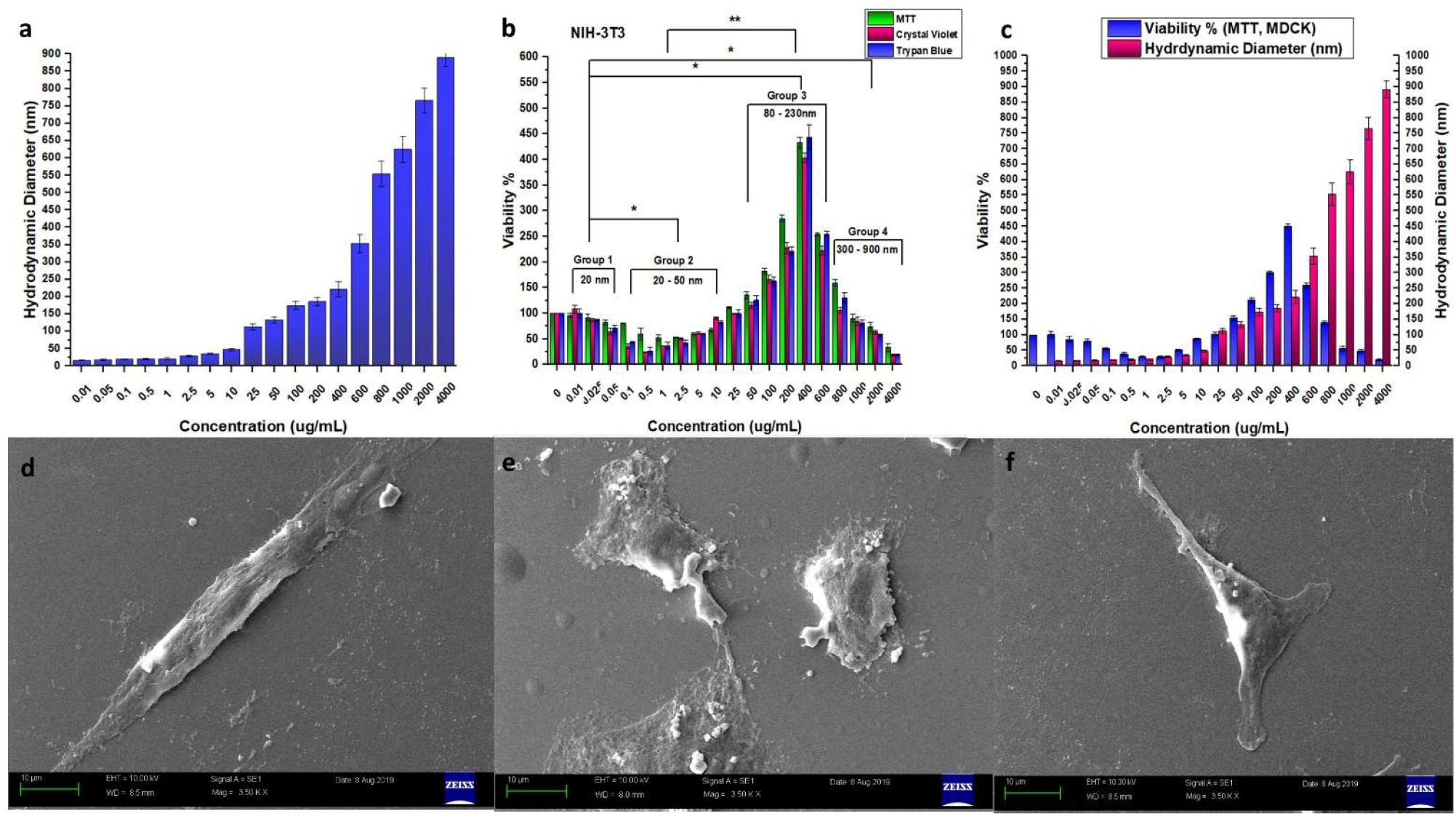
(a) Hydrodynamic diameter of Fullerene nanoparticles obtained by DLS in the concentration range from 0.01 μg/mL to 4000 μg/mL. The size remained almost same from 0.01 to 1 μg/mL and then increased gradually up to 400 μg/mL. Above this concentration (400 μg/mL), the size increases drastically. (b & c) Cell viability trend of NIH-3T3 with (b) original concentration of Fullerene used and (c) size of the nanoparticles formed with each concentration used. The cell viability is lower than the control for 20-50 nm sized particles whereas viability increases for 130-230 nm sized particles. (d-f) SEM, of MDCK cells treated with (a) only media, (b) F_T30_, (c) F_P200_. F_T30_ treated cells show altered morphology; Control and F_P200_ treated cells show normal morphology.

Therefore, we considered Fullerene nanoformulation prepared in the DMEM+FBS system for further studies. We measured the hydrodynamic diameter of the particles using dynamic light scattering (DLS) measurement (**†ESI S1**). The z-average diameter of the dispersed particles in the nanoformulation remained ~20 nm for the concentration range from 0.01 to 1 μg/mL and then increased gradually from 20 nm (1 μg/mL) to ~230 nm (400 μg/mL) (**†ESI Table S1**). The hydrodynamic diameter of the particles above 400 μg/mL increased drastically from 350 nm (600 μg/mL) to ~900 nm (4000 μg/mL) (**Figure 1 a**). The low polydispersity index characterized by a value of 0.11±0.03 was obtained in the concentration range of 0.01 to 400 μg/mL. At this range, the zeta potential was found to be −18±5 millivolts. This signifies repulsion among the particles resulting in a moderately stable suspension, suggesting their suitability in biomedical applications. Size distribution analysis by number % showed a similar size as z-average diameter (**†ESI Fig S1**). Above this concentration (400 μg/mL), the dispersion was poor and was characterized by rapid settling of the particles, low zeta potential (−8±4 millivolts.) and, a high polydispersity index of 0.56±0.07.

We used 6 cell lines of multiple tissue origins (MDCK, NIH-3T3, A549, HepG2, SH-SY5Y, and BMSC) for evaluating the cell viability activity of the nanoformulation. The cells were cultured in DMEM+10%FBS media. To check the effect of nanoformulation on cells, 4 orthogonal tests were performed. MTT, Crystal violet, Trypan blue, and LDH assay after 24 hours’ treatment (**†ESI S2**). In all the viability assays, 4 trends were obtained (**Figure 1 b**). In the first group (**Group 1: 0.01-0.05 μg/mL, ~20 nm**), no apparent activity was obtained. In the second group (**Group 2: 0.1-10 μg/mL, ~20-50 nm**), cytotoxicity was observed. In the range of 0.1 μg/mL to 1 μg/mL, a gradual increase in cytotoxicity was seen. DLS showed a constant size of ~20 nm in this concentration range, suggesting that 20 nm particles are toxic. However, as the size increased up to 50 nm, the toxicity started decreasing. In the third group (**Group 3: 25-400 μg/mL, ~80-230 nm**) cell viability increased. The viability reached ~4-fold in some cell lines (MDCK, NIH-3T3, A549, and HepG2) and 2-3-fold in others (SH-SY5Y, BMSC) (**†ESI Fig S2 a-e**). We observed maximum proliferation at 200±30 nm particle size obtained from 400 μg/mL Fullerene dispersion. Above this concentration (**Group 4: 600 to 4000 μg/mL, ~300-900 nm**), cytotoxicity was again observed. LDH cytotoxicity tests showed that the nanoformulation made from 5 μg/mL (~35 nm) and 10 μg/mL (~50 nm) concentrations show lower cell viability and increased cytotoxicity in LDH assay. On the other hand, the concentrations showing neutral or proliferative action on cells (25 μg/mL (~90 nm); 50 μg/mL (~120 nm); 200 μg/mL (~150 nm) and 400 μg/mL (~220 nm)) did not show any significant cytotoxicity effect on the mammalian cells in LDH assay (**†ESI Fig S2 f**). To corroborate this data, fluorescence microscopy and scanning electron microscopy analysis were carried on one representative nanoparticle size from each group: (Toxic; 30 nm; FT30) and (Proliferative; 200 nm; FP200) (†ESI S2). SEM analysis of MDCK cells showed that while there were no morphological changes in control and FP200 treated cells (Figure 1 d, f), FT30 treated cells showed altered morphology (Figure 1 e). DAPI (4‘,6-diamidino-2-phenylindole) and FDA staining also confirmed that in the control and FP200 treated cells, there were no abnormal nuclei and more number of viable cells, whereas FT30 treated cells showed fragmented nuclei and lower number of viable cells (†ESI Fig S3) (ImageJ; Par8cle analysis).

Next, we wanted to test the effect of different amounts of the prepared nanoparticle sizes on cell viability. (†ESI Fig S4 a,b) For this, we first prepared 30, 50 and 80 nm sized particles. MDCK cells were treated with each of these prepared nanoparticle with 1, 10 and 100 μg/mL nanoparticle concentrations. 30 nm particles showed cytotoxicity which increased with concentration. For 50 nm, the toxicity observed at each amount was found to be lower than 30 nm. For 80 nm sized particles, similar viability was observed as that of control. Next, we prepared 150, 170 and 200 nm sized particles. A stock solution of 200 μg/mL was prepared for each sized particles. To prepare stock solution, the initial suspension was either concentrated or diluted as per requirement to obtain final concentration of 1, 10 and 100 μg/ml. After dilution/concentration, the size of the particles did not change significantly, as characterized by DLS (**ESI Section S2: Size vs Concentration assay**) These were then applied on cells. None of them were effective at 1 μg/ml concentration suggesting that this amount is too low for these particles to exert any effect. The 150, 170 nm and 200 nm sized particles showed an enhanced cell proliferation at 10 and 100 μg/ml. The 200 nm sized particles increased cell viability to a greater extent as compared to the 150 and 170 nm particles. The results confirm that the size of the nanoparticles formed by dispersion of different amounts of fullerene play the primary role in determining the biological activity and increased amount of nanoparticles augments the biological activity.

We next evaluated the biological effect of Fullerene nanoformulation on 3D spheroids formed from MDCK and HepG2 cell lines due to its ease of forming spheroids. 3D culture is more appreciated for drug evaluation due to their relevance to *in-vivo* structures. In 3D culture of both cell lines, the proliferation effect (F_P200_) and cell death (F_T30_) were 2-fold more than the control in 24 hours (**†ESI Fig S4 c, d**).

Since F_P200_ caused cell proliferation, it was imperative to check whether the treated cells were transformed into invasive cells, a feature of neoplasia. Migration assays showed that F_P200_ treated cells (MDCK: non-cancerous and HepG2: liver cancer cell line) did not migrate through a 200 nm pore size membrane filter and behaved like control (**†ESI Fig S4 e, f**).

In order to translate the proliferative effect of Fullerene nanoformulation (F_P200_) into a suitable biomedical application, we hypothesized that this effect might be used in tissue repair. The lengthiest phase of wound healing i.e. phase 3 involves fibroblast proliferation followed by epithelial cell proliferation in phase 4. Thus, we used cellular and mice wound models (**Figure 2**) to check tissue repair. Since the proliferative particles increase epithelial (MDCK) and fibroblast (NIH-3T3) proliferation (**Figure 1 b, †ESI Fig S2 d**), a scratch assay on cell monolayers was employed. The wound closure was checked after every 4 hours by FDA staining of the cells (**†ESI S3**). The control experiments (**Figure 2 a-c**) showed a gradual wound closure but the wound required 36 hours to close fully.

**Figure 2:**
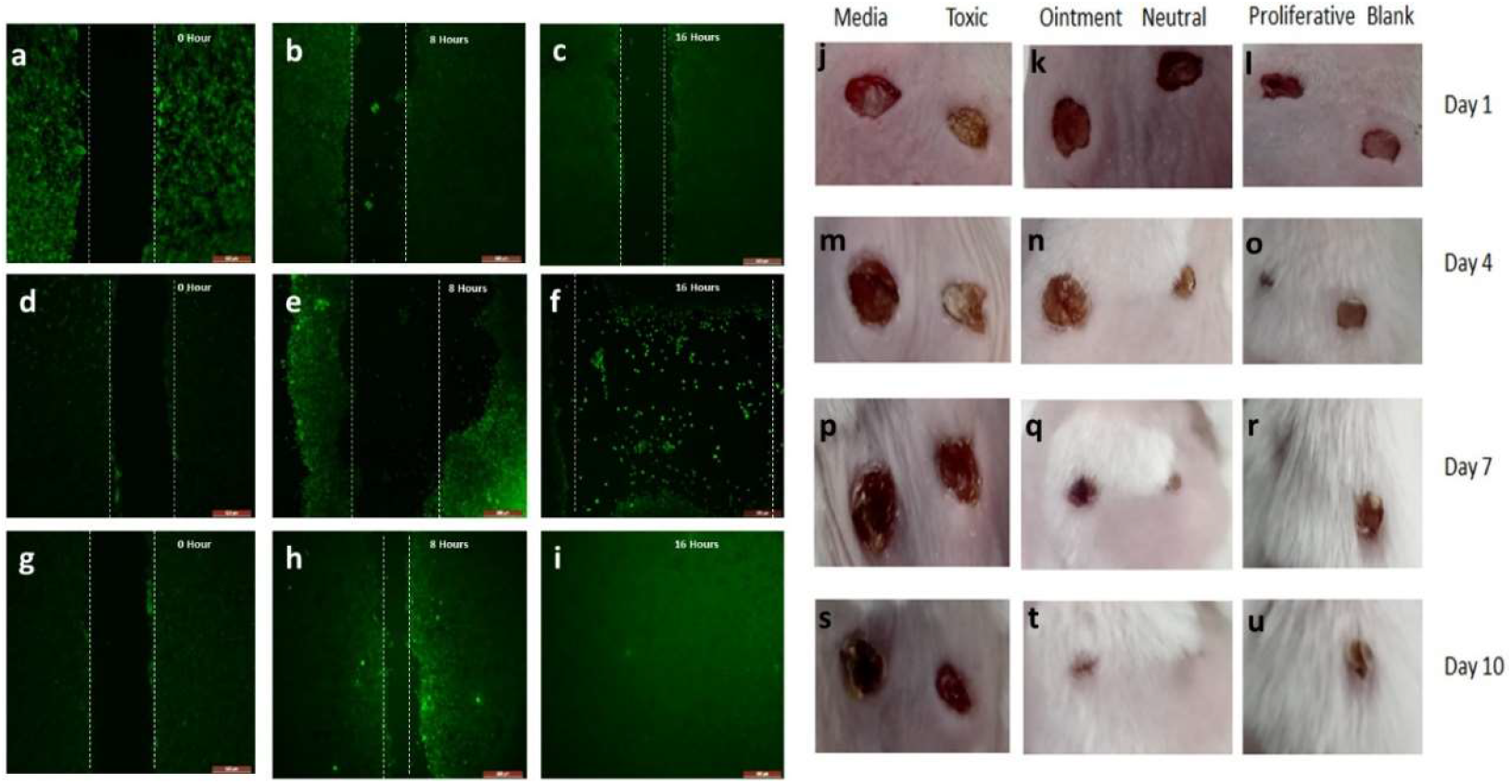
(Left Panel) Scratch assay on NIH-3T3 cells, normal media. (Scale bar 200 μm). Control cells without treatment (a-c) show slow wound closure and incomplete recovery in 16 hours; FT30 treated cells (d-f) show gradual expansion of the wound area; F_P200_ treated cells (g-i) show accelerated wound closure and complete repair within 16 hours. (Right Panel) Circular (5 mm diameter) full-thickness wounds showing healing progress throughout 10 days. F_P200_ (proliferative) nanoformulation treated wounds (l,o,r,u) heal at the fastest rate with accelerated hair growth. F_N80_ (Neutral) and ointment treatment (k,n,q,t) shows an intermediate rate of wound closure, slower than proliferative but faster than the control experiments (Media, Blank,) and F_T30_ (Toxic) (j,m,p,s) which show the slowest rate.

In F_T30_ treated cells (**Figure 2 d-f**), instead of wound closure, wound area expanded due to increased cell death. Interestingly, F_P200_ treated cells showed an accelerated wound closure, and complete repair was seen within 16 hours (**Figure 2 g-i**). Quantification of the wound area by ImageJ showed that the control and F_N80_ nanoformulation (80-100 nm size, 25-50 μg/mL; no cell viability effect; **Figure 1 b**) led to the wound closure in the same pace. In F_T30_, the wound area % increased with time in contrast to F_P200_ where wound area closed at an accelerated pace. Besides acute healing, accelerated wound care is a requisite in delayed healing conditions such as diabetes mellitus.

When the *in-vitro* scratch assay was repeated in hyperglycemic conditions, the overall wound healing was seen to be slower as compared to normal media but a similar trend followed. Control and F_N80_ treated cells showed complete recovery in 40 hours compared to the F_P200_ accelerated effect where the wound was repaired in 30 hours. The F_T30_ showed wound area expansion throughout 40 hours (**†ESI Fig S5**).

One of the reasons for the failure of nanomedicine translation is the non-reproducibility of *in-vitro* results in *in-vivo* animal models. For successful design-based biomedical application of the proliferative nanoformulation, circular full-thickness wounds 5 mm were created on BALB/c mouse strain. For each group, 2 mice were taken and the number of wounds checked for each set was 4. All the animal experiments were approved by the IAEC committee. The mice wounds were monitored every alternate day and quantified using Vernier caliper (**#x2020;ESI Fig S6**). After 14 days of complete healing, the healed tissues were excised and Hematoxylin & Eosin (H&E) staining was done to check tissue integrity (**†ESI S4**). **Figure 2 (j-u)** shows the skin tissue repair of the mice. In both media and F_T30_ (cytotoxic nanoformulation) treated wounds (**Figure 2 j,m,p,s**), the healing was very slow. The F_N80_ (neutral) nanoformulation and the ointment Mupirocin (**Figure 2 k,n,q,t**) showed almost similar healing activity. The F_P200_ (proliferative) however showed the most promising effect obvious from day 4 and the whole wound was healed with regrowth of hair within 10 days (**Figure 2 l,o,r,u**). (H&E) staining showed that the control experiments have a similar thickness of the epidermis, a moderate number of blood vessels, and sebaceous glands as visualized and quantified by ImageJ. F_P200_ treated wounds show similar morphology except that the number of hair follicles and sebaceous glands are more in number (ImageJ quantification and scoring). On the other hand, F_T30_ treated wounds show a poor tissue deposition (**†ESI Fig S7**). Another 14 days were devoted for checking any side-effects on the animals but no effects were apparent.

We have prepared a dispersion system for pristine Fullerene in routine cell culture media. The resulting formulation exhibits concentration-based formation of discrete sized nanoparticles. Being made in a biologically relevant media, the formulation obtained can be used for multiple applications where proliferation or reduction in cell numbers are required. Lack of organic solvents or technically intricate steps in the formulation process leads to better feasibility for clinical translation. Observation of the toxic and proliferative effects obtained by the nanoformulation may not be generalized for Fullerene dispersed by other methods or raw material composition. Deciphering of comprehensive understanding of these mechanisms of these effects in future will help in understanding them. However, Fullerene and its derivatives have been already implicated in multiple biological pathways such as antioxidant or pro-oxidant effects and cell death pathways (15) and in case of our nanoformulation also, these pathways might be the major key factors.

Considering proliferative effect as a rationale for size range of 130-230 nm particles of Fullerene, we have shown one application for successful skin wound healing. The quality attribute and this rationale approach followed here will provide a way to use nanoparticles in general when biomedical application is the ultimate goal. Besides skin wound-healing, the proliferative nanoformulation may also be able to find multiple applications as regenerative medicine for different tissues/organs.

## Supporting information

Electronic Supplementary Information

## References

1 H. F. Krug, Angewandte Chemie International Edition, 2014, 53, 12304–12319.

2 K. Greish, Methods in molecular biology (Clifton, N.J.), 2010, 624, 25–37.

3 L. Wang, C. Hu and L. Shao, International journal of nanomedicine, 2017, 12, 1227–1249.

4 P. Sharma, S. Brown, G. Walter, S. Santra and B. Moudgil, Advances in Colloid and Interface Science, 2006, 123-126, 471–485.

5 M. Fathi-Achachelouei, H. Knopf-Marques, C. E. Ribeiro da Silva, J. Barthès, E. Bat, A. Tezcaner and N. E. Vrana, Front Bioeng Biotechnol, 2019, 7, 113–113.

6 N. Gharbi, M. Pressac, M. Hadchouel, H. Szwarc, S. R. Wilson and F. Moussa, Nano Letters, 2005, 5, 2578–2585.

7 S. Kato, H. Taira, H. Aoshima, and Y. Saitoh, et al. “Clinical Evaluation of Fullerene-C(60) Dissolved in Squalane for Anti-Wrinkle Cosmetics.” Journal of nanoscience and nanotechnology, 2010, 10: 6769–6774

8 Kovochich M, Espinasse B, Auffan M, et al. Environ Sci Technol. 2009;43(16):6378–6384.

9 J. Kolosnjaj, H. Szwarc and F. Moussa, Bio-Applications of Nanoparticles, ed. W. C. W. Chan, Springer New York, New York, NY, 2007, 13, pp. 168–180.

10 T. Baati, F. Bourasset, N. Gharbi, L. Njim, M. Abderrabba, A. Kerkeni, H. Szwarc and F. Moussa, Biomaterials, 2012, 33, 4936–4946.

11 J. K. Folkmann, L. Risom, N. R. Jacobsen, H. Wallin, S. Loft and P. Møller, Environmental health perspectives, 2009, 117, 703–708.

12 Ekaterina S. Kovel, Anna S. Sachkova, Natalia G. Vnukovia, Grigoriy N. Churilov, Elena M. Knyazeva and Nadezhda S. Kudryasheva. Int J Mol Sci, 2019 May; 20(9):2324.

13 G. V., Andrievsky, Marina. Kosevich, Oleh. Vovok, Vadim Shelkovsky and Lyudmila Vashchenko Journal of the Chemical Society, Chemical Communications 1995 (12): 1281–1282.

14 Koichi Imai, Fumio Watari, Tetsunari Nishikawa, Akio Tanaka, Akito Tanoue, Kazuaki Nakamura and Hiromasa Takashima. Nano Biomedicine, 2011, 3(2), 288–293.

15 Orlova A.M., Trofimova T.P., Shatalog O.A, Der Pharmacia Lettre, 2013, 5 (3), 99–139.

