## Supplementary material for "Accelerated tissue repair through cell proliferative effect by a size controlled aqueous based Fullerene C_60_ nanoformulation": Electronic Supplementary Information

a	Nabodita Sinha
Department of Biological Sciences and Bioengineering
Indian Institute of Technology Kanpur, Uttar Pradesh – 208016, India.

b	Avinash Y.Gahane

Department of Biological Sciences and Bioengineering
 Indian Institute of Technology Kanpur, Uttar Pradesh – 208016, India

C	Professor Ashwani Kumar Thakur
 Department of Biological Sciences and Bioengineering

Indian Institute of Technology Kanpur, Uttar Pradesh – 208016, India.


**Table of Contents:**

| **S No** | **Name** | **Title** | **Page** |
| --- | --- | --- | --- |
| **1** |  | **Instrument Details** | **1** |
| **2** | **S1** | **Preparation and optimization of Fullerene dispersion in aqueous media and their characterization** | **2** |
| **3** | **S2** | ***In-vitro* biological activity of the nanoformulation on 2D and 3D cell cultures** | **3** |
| **4** | **S3** | **Scratch assay in normal and hyperglycemic media** | **6** |
| **5** | **S4** | ***In-vivo* wound healing assay** | **7** |
| **6** | **Table S1** | **Hydrodynamic diameter of Fullerene nanoformulation in DMEM+FBS with concentration** | **9** |
| **7** | **Figure S1** | **Size distribution analysis of nanoparticles** | **10** |
| **8** | **Figure S2** | **Cell viability assay on different cell lines** | **11** |
| **9** | **Figure S3** | **Fluorescence staining for nuclear morphology and viability of cells** | **12** |
| **10** | **Figure S4** | **3D Cell culture, size vs. concentration and migration assays** | **13** |
| **11** | **Figure S5** | ***In-vitro* scratch assay quantification** | **14** |
| **12** | **Figure S6** | ***In-vivo* wound repair quantification** | **15** |
| **13** | **Figure S7** | **Hematoxylin Eosin staining of excised wound tissue** | **16** |

**Instrument Details:**

The following instrument models were used for the work:

Malvern Zetasizer Nano ZS90 (633 nm laser, scattering angle 90^0^) for DLS measurements such as hydrodynamic diameter, polydispersity index and zeta potential; Leica DM 2500 for bright field and fluorescence microscopy; Perkin Elmer ELISA Microplate Reader for all absorbance assays in microplates; Carl Zeiss EVO18 for Scanning Electron Microscopy of cell morphology; Leica Ultramicrotome and Embedding systems for tissue processing and sectioning.

**Detailed Experimental Methods:**

**Section S1. Preparation and optimization of Fullerene dispersion in aqueous media and their characterization:**

Fullerene C_60_ powder (98% pure) was obtained from Sigma Aldrich. Four aqueous media i.e. double distilled water, phosphate buffer saline (PBS), Dulbecco Modified Eagle Medium (DMEM) and DMEM+10% FBS (Fetal Bovine Serum), were selected to observe their suitability as dispersion media for Fullerene. Four representative concentrations were at first chosen to test the dispersion method: 0.1, 10, 100 and 1000 µg/mL. For preparing the lower concentrations i.e. 0.1 and 10 µg/mL, 1 L glass bottles and for preparing the higher two concentrations, 10 mL glass vials were used to ensure accurate weighing and efficient use of the raw material. Fullerene powder was then accurately weighed and added to the bottles/vials.

| Fullerene powder amount (mg) | Bottle/Vial volume  (mL) | Media to be added  (mL) | Effective concentration (µg/mL) |
| --- | --- | --- | --- |
| 0.1 | 1000 | 1000 | 0.1 |
| 10 | 1000 | 1000 | 10 |
| 1 | 10 | 10 | 100 |
| 10 | 10 | 10 | 1000 |

The vials with powder were then autoclaved for sterilization at 121^0^C, 15 minutes. The 4 types of media (water, phosphate buffer saline, DMEM and DMEM+10% FBS) were then sterile filtered individually through 200 nm filter membranes and added to the vials at required amount under aseptic conditions. The mixtures were then kept for stirring on a magnetic stirrer at 700 rpm (rotations per minute) and 25^°^C for 14 days. Different speeds (300, 500, 700, 900 rpm) and different time durations (1-21 days) were checked for process optimization for each type of media and each concentration.

Dyamic Light Scattering (DLS) method was used to measure the hydrodynamic diameter and zeta potential of the formed particles. For each of the concentrations mentioned above, the DLS measurement was performed every 24 hours for 21 days at each stirring speed. For both size and zeta potential measurement, 1 mL of sample was used. Parameters such as Z-average, polydispersity index and zeta potential was recorded for each sample in one of the four media systems. Considering the parameters such as discrete sizes, low polydispersity index and high zeta potential, the following conditions were subsequently chosen for optimum dispersion: DMEM+FBS as media, 700 rpm and 14 days as reaction conditions.

DMEM+FBS was then used to prepare Fullerene concentrations from 0.01 µg/mL to 4000 µg/mL by stirring for 14 days at 700 rpm at 25^°^C. For each of these concentrations, the Z-average, polydispersity index and zeta potential were recorded. The size distribution analysis was done for each concentration using number weighing method after 14 days.

**Section S2.** ***In-vitro* biological activity of the nanoformulation on 2D and 3D cell cultures**:

**Cell culture protocols:**

The cell lines used in the study are: MDCK (canine kidney epithelial), NIH-3T3 (mouse skin fibroblast), A549 (human lungs epithelial), HepG2 (human liver epithelial), SH-SY5Y (human neuroblastoma) and BMSC (mouse bone marrow stromal cells). All the cell lines were cultured in DMEM+10% FBS media. The frozen vials of the cells were thawed at 37^0^C in a water-bath and centrifuged at 2000 rpm for 1 minute. The supernatant was discarded and the pellet was resuspended in 1 mL of fresh media. The cell suspension was then added to T25 flasks along with 4 mL of fresh media and incubated at 37^°^C, 10% CO_2_ until the cells reached 80% confluence. Once cells reached desired confluency, the media was discarded and the cells were trypsinized by adding 1 mL of 0.25 % Trypsin+0.02% EDTA except SH-SY5Y for which 0.05% Trypsin was used. The flasks were kept in the incubator at 37^°^C for 10 minutes to allow cell detaching from flask surface. To the detached cells, 2 mL of fresh media was added to neutralize trypsin. To 90 µL of the cell suspension, 10 µL of 0.4% of trypan blue was added and mixed well. The stained suspension was then applied on a hemocytometer and bright-field microscopy was used for counting the number of live vs. dead cells under 10X objective. The dead cells take up trypan blue and thus appear blue while the live cells appear colourless with discernible cell border. The average number of live and dead cells were counted for each of the large squares in the set of 16 squares in the hemocytometer. The formula used for calculating viability and cell density is as follows:

**Viability: (Average Live cell count) / (Average Dead cell count)**

**Cell Density (no. of cells/mL) = (Average no. of viable cells in each square) ×10000×Dilution factor**

The cells were then either used for further passaging or freezing or in biological assays as described below.

To check the effect of the nanoformulation on cells, 4 orthogonal tests were performed. **MTT**(3-(4,5-dimethylthiazol-2-yl)-2,5-diphenyl tetrazolium bromide; Sigma Aldrich): for mitochondrial metabolism, **Crystal violet** (Sigma Aldrich): for cellular biomass, **Trypan blue** (Merck): for viability and **LDH** (Lactate dehydrogenase; Pierce-Thermofisher) assay: for cytotoxicity. For each of the assays, the cells were trypsinized and the cell concentration was calculated as described above. 10000 cells were then seeded per mL into 96-well assay plates (except trypan blue assay where 24-well plates were used). To the cells, nanoformulation of different concentrations (0.01 to 4000 µg/mL; size ranging from 20 to 800 nm) were added. For 96-well plates, 100 µL cell suspension was treated with 100 µL of the nanoformulation; for 24-well plates, 500 µL cell suspension was treated with 500 µL nanoformulation. After treatment, the cells were incubated for 24 hours at 37^°^C, 10% CO_2_. After 24 hours, the old media was aspirated and the cells were washed twice with PBS.

In all the assays, besides control experiment, blank sets were also implemented without cells to check for any interference of the assay reading due to nanoformulation alone. The following assays were performed:

**MTT Assay**:

0.5 mg/mL MTT solution was prepared in DMEM+10% FBS. 20 μL of MTT was added to each well (cell density of 10000 cells/mL) and the cells were incubated for 2 hours at 37^°^C Centigrade. To the wells, 50 μL of DMSO (Fisher Scientific) was added to lyse the cells and absorbance was checked at 530 nm using Perkin Elmer ELISA microplate reader.

**Crystal Violet Assay**:

Crystal violet stain was prepared by dissolving 0.2 g in 2% ethanol (100 ml). 50 μL of crystal violet was added to each well (cell density of 10000 cells/mL) and incubated for 10 minutes at room temperature. The wells were washed twice with PBS to remove dead and detached cells. The stain was then solubilized with 1% SDS and plate was agitated on shaker for 10 minutes. Absorbance was then measured at 570 nm using microplate reader.

**Trypan blue Assay:**

The cells were trypsinized as mentioned above. 0.4 % trypan blue (10 µL) solution was added to 90µL of cell suspension (cell density of 10000 cells/mL) and the viability was calculated according to the method described above by hemocytometer (**Section S2: Cell Culture Protocols).**

**LDH Assay:**

For LDH assay, the following concentrations of the nanoformulation were used: 5,10, 25, 50, 200 and 400 µg/mL. After 24 hours of treatment, 50 μL of supernatant from the treated cells, were added to fresh 96-well plates since the LDH enzyme to be assayed is released into the extracellular media. The extent of LDH release depends on the cellular membrane integrity. The more the membrane integrity is hampered due to cytotoxicity, the more LDH is released. 50 μL of reaction mixture containing lactate, diaphorase and NAD^+^ were added to the samples which act as the substrate and cofactors for the reaction. The mixture was incubated at room temperature for 30 minutes. The absorbance of the red formazan product so formed was measured at 490 nm using microplate reader which is directly proportional to the amount of LDH released by the cells and hence gives a measure of the amount of cytotoxicity.

**Choosing representative concentrations and sizes for further assays:**

From the interpretation of the results of the biological assays, we obtained biological effects based on size (**Manuscript Page 2**), DLS and biological assays showed that the size of 20 to 50 nm (0.1 µg/mL to 10 µg/mL) are cytotoxic, 80-100 nm (25 µg/mL to 50 µg/mL) are neutral, 130-230 nm (100 µg/mL to 400 µg/mL) are proliferative. Further experiments were performed with the representatives of these size groups: F_T30_ and F_T50_ (Cytotoxic; 30 and 50 nm diameter respectively), F_N80_ (Neutral i.e. no apparent biological effect on cells; 80 nm diameter), F_P150_ and F_P200_ (Proliferative; 150 nm and 200 nm diameter).

**3D Cell culture and viability assay:**

3D cell culture requires low cell adhesion to assay plates. For this purpose, 1.25% of agarose solution was prepared in PBS, sterilized and 60 µL of this solution was added to each well of a fresh 96-well plate. After the agarose coated wells solidify, 10000 cells (50 µL) of either MDCK or HepG2 cell lines were seeded on each of these wells. The cells were then treated with 50 µL of F_T30_ (Cytotoxic, 30 nm, 1 µg/mL), F_T50_ (Cytotoxic, 50 nm, 10 µg/mL), F_N80_ (Neutral, 80 nm, 25 µg/mL), F_P150_ (Proliferative, 150 nm, 200 µg/mL) and F_P200_ (Proliferative, 200 nm, 400 µg/mL). The cells were incubated for 4 days till 3D spheroids were formed. 3D spheroids were then dissociated by pipetting. MTT and trypan blue assays were then performed to check cell viability.

For all the above assays, each concentration was applied to cells in at least 9 wells; therefore, technical replicates per concentration were 9. Each of these assays were performed on 3 different days; therefore, biological replicates were 3. Thus for each concentration, total replicates were 9×3 = 27.

**Visualization of nanoformulation effect on cells by microscopy:**

For these experiments: cells were trypsinized, counted by hemocytometer and 10000 cells/mL were seeded on 12 mm coverslips in 24-well plates. To 500 µL of the cells, 500 µL nanoformulation was added. One set was control experiments where no treatment was added. To the other two sets, F_T30_ (1 µg/mL) and F_P200_ (400 µg/mL) were added respectively.

**SEM (Scanning electron microscopy):**

12 mm coverslips were sterilized with 70% ethanol and 1 hour UV duration and placed in 24-well plate. After 24 hours, coverslips were washed with PBS and cells were fixed with 2.5% glutaraldehyde. The coverslips were then subjected to ethanol gradient dehydration (25%, 40%, 60%, 80%, 90% and 100%) and then dried in a vacuum desiccator for 6 hours. The coverslips were then attached on SEM copper grids with conductive carbon tape, sputtered with gold and visualized in SEM.

**Fluorescent Microsocopy:**

**DAPI (4^‘^,6-diamidino-2-phenylindole) Staining:**

After 24 hours of nanoformulation treatment, the cells were washed twice with PBS. These are then fixed with 2.5% glutaraldehyde for 10 minutes. The fixing agent was then washed away with PBS and permeabilizing agent 0.2% Triton X was added. The cells were kept at room temperature for 10 minutes and 300 nano-Molar DAPI in PBS is added to it. The cells are kept in dark for 10 minutes and visualized under fluorescence microscopy for observing micronuclei and DNA condensation.

**Viability staining by FDA:**

FDA (Fluorescein diacetate) staining solution was prepared at a concentration of 0.5 mg/mL in acetone. The FDA staining solution was prepared by adding 8 μL FDA to 5 mL PBS. The cells were washed twice with PBS and 20 μL of the staining solution is added. The samples were then visualized under fluorescent microscope.

The above images were analysed in ImageJ. For counting the number of viable cells in FDA, the Analyze and Measure features of ImageJ software were used and corroborated with manual counting. For each type of treatment, 3 different areas on slides for each sample were counted.

**Migration Assay:**

MDCK and HepG2 cell lines were used in this assay. Corning 200 nm transwell filters were placed on 24-well plates after adding 500 µL fresh media to the lower chamber. The filter was then seeded with 10000 cells/mL of either MDCK or HepG2 cells. Three sets of experiments were used. In 1^st^ set, the cells were treated with the nanoformulation (F_T30_, F_T50_, F_N80_, F_P150_ and F_P200_) after seeding on the filter. In this set its assumed that although the cells cannot diffuse through the membrane pores, the Fullerene particles can diffuse freely and can establish a concentration gradient. Thus it may be observed whether in response to the established concentration gradient, the cells migrate through the pores or not. In the 2^nd^ set, the cells were first treated with the respective nanoformulation treatment and then seeded on the filters. Thus the lower chamber had only media, but the upper chamber had treated cells. This set was performed to observe whether the treated cells migrate after coming in contact with normal media to check for transformed cells. In the 3^rd^ set, the lower chamber had media along with nanoformulation but the cells were not directly treated. The cells were exposed to the particles only through media contact. This set was to check the effect of the nanoformulation as a chemotactic agent on the cells i.e. whether the cells could migrate through the pores by sensing these particles. After 24 hours of the experiment, the transwell filters were isolated from the 24-well plates and the cells from the apical part of the filters were scraped out using cotton swabs. The filters were then inverted and stained with 0.2% crystal violet stain for 10 minutes as already described above. The filters were washed repeatedly in double distilled water to remove the excess stain and then dipped into 500 µL of 1% SDS for dissolving crystal violet by agitating on a shaker for 10 minutes. The filters were then removed and the amount of stain is measured by taking the absorbance at 570 nm using a microplate reader.

**Size vs. Concentration Assay:**

First we prepared 30 nm by dispersing 2.5 µg Fullerene per mL media; 50 nm by dispersing 10 µg per mL media and 80 nm nanoparticles by dispersing 25 µg Fullerene per mL media according to our developed method by stirring for 14 days. These nanoparticles attain the sizes of 30, 50 and 80 nm respectively after effective dispersion. Next we prepared 3 types of nanoparticles which show proliferation effects. 150 nm particles were prepared by dispersing 100 µg per mL media, 170 nm by dispersing 200 µg per mL media and 200 nm by dispersing 400 µg per mL media. A stock solution of 200 µg/mL was prepared for each size. For the smaller particles, since the initial concentration of preparation was low, the prepared nanoparticles were centrifuged and the small volume of media was added to the pellets to increase their concentration. This did not alter the physicochemical properties of the particles. All the smaller sizes were initially prepared at a volume of 100 ml. This was then divided into two centrifuge tubes (50 ml each). The tubes were centrifuged at 6000 rpm for 20 minutes. Although varying centrifugation speeds were employed for this process, the conditions of 6000 rpm and 20 minutes was found to be optimum without compromising the stability of the particles. The supernatant was discarded and to the pellet in two flasks, the required final volume of fresh media was added to obtain the final concentration. The suspension was mixed gently with a 1 ml pipette and transferred to fresh sterilized vials. For 200 µg /ml, no such steps were required. For 400 µg/ml, a simple dilution was made for converting it to 200 µg/ml by adding double volume of media. 1 ml of the sample was withdrawn then for DLS characterization.

To further confirm that the final concentration is according to our desired value, a toluene extraction method was employed to calculate the actual concentration. For this, 10 ml of the sample was extracted into equal volume of toluene sample by stirring at 700 rpm for 30 minutes. The organic layer was then extracted and evaporated to half the volume using vacuum desiccator for 5 minutes. The samples were then checked for absorbance at 336 nm for Fullerene absorbance. The concentration was calculated by using a standard curve of Fullerene standards. The following table further specifies each and every step followed for characterization. This way of presentation will further provide answer to the reviewer question.

Table:

| Initial Amount  µg | Initial Volume  ml | Initial Size  nm | Centrifugation Volume and amount | Total Media Volume added  ml | Final theoretical concentration  µg/ml | Final experimental concentration  µg/ml | Final Size by DLS  nm |
| --- | --- | --- | --- | --- | --- | --- | --- |
| 2.5 | 1 | 28.16 | 250 µg in 100 ml | 1.25 | 200 | 194.75 | 29.24 |
| 10 | 1 | 47.26 | 1000 µg in 100 ml | 5 | 200 | 195.78 | 53.45 |
| 25 | 1 | 82.5 | 2500 µg in 100 ml | 12.5 | 200 | 194.91 | 86.98 |
| 100 | 1 | 153.7 | 10000 µg in 100 ml | 50 | 200 | 196.21 | 156.81 |
| 200 | 1 | 175.4 | Not required | - | - | - | - |
| 400 | 1 | 221.1 | - | 1 | 200 | 199.75 | 211.7 |

These prepared nanoparticles having the desired size were then applied on cells at 3 concentrations i.e. 1, 10 and 100 µg/mL. Each of these prepared nanoparticles were applied on cells at 3 concentrations: 1, 10 and 100 µg/ml i.e. the cells within a group were getting similar size of nanoparticles but the amount of these nanoparticles available to the cells varied. The MDCK cells were trypsinized and seeded on 96-well plates as described before (Section S2; **Cell culture protocols**) at a density of 10000 cells/mL. To 100 µL of cell suspension, 100 µL of the nanoformulation was added, making the effective concentrations of the nanoformulation available to the cells as: 1, 10 and 100 µg/mL. The cells were then incubated for 24 hours and MTT assay was done to check viability of the cells.

**Section S3: Scratch Assay in Normal and Hyperglycemic media**

MDCK and NIH-3T3 cells were seeded at a density of 20000 cells/mL on 4 well plates and allowed to form a monolayer. With a 1 ml pipette tip, a scratch was made on the monolayer vertically to simulate a wound. The media was then aspirated to remove the dead and detached cells. The cells were washed twice with PBS and fresh media (500 µL) and nanoformulation F_T30_ (10 µg/mL) or F_P200_ (400 µg/mL) (500 µL) were added. The wound closure was then checked after every 4 hours by FDA staining the cells which enables clear visualization of the wound gap.

In another set of experiments, hyperglycemic media was used to simulate the high-glucose conditions of a diabetic patient. The same assay was repeated but the media was supplemented with dextrose at a concentration of 3 mg/mL. The available dextrose thus available to the cells was 1.5 mg/mL, simulating diabetic conditions.

**Section S4**: ***In-vivo* wound healing assay**

All the animal experiments were carried out under aseptic conditions. The mice were kept at cages throughout their lifetime with 2-3 animals per cage. The cages were individually ventilated with continuous air flow. They were kept at designated institute animal facility with 12h:12h light/dark cycle. Food and water was provided to them *ad libitum*. For creating wounds, BALB/c mice were anaesthetized with 0.1 ml ketamine per animal. The dorsal hairs were shaved by an electric razor and depilating cream. Identical wounds of 5 mm were created on both sides of the backbone by using biopsy punch. Appropriate treatments (Media, Mupirocin ointment, toxic (F_T30_), neutral (F_N80_) and proliferative (F_P200_) were then applied on the wounds (20 µL, containing 0.008 µg nanoformulation). Diameter of wounds were measured on alternate days with vernier caliper. After 14 days of complete healing, the healed tissues were excisioned with surgery scissors after anaesthesizing the mice and Hematoxylin & Eosin (H&E) staining was done to check tissue integrity and healing parameters.

**Hematoxylin and Eosin Staining:**

Tissue Fixation: The excised tissues were immediately drop-fixed in 4% neutral buffered formalin solution (2 ml formalin per 100 g of tissue). The tissues were fixed for 24 hours at room temperature. The fixed tissues were then used immediately for processing.

Tissue processing: The fixed tissues were dipped into the coplin jars and following steps were followed:

- 70% ethanol for 30 minutes.
- 80% ethanol for 30 minutes.
- 95% ethanol for 30 minutes.
- Absolute alcohol for 1 hour.
- Xylene for 1 hour.
- Paraffin wax at 60^0^C for 1 hour

**Embedding and sectioning of tissues:** A small amount of molten paraffin was poured into molds with cassette in place and the tissue was transferred to it with the help of forceps placing the cut side down. More paraffin was added to the tissue to fill up the mold and these were transferred to cold plate (60^0^C) to allow the paraffin to solidify. After 30 minutes, the tissue block with the cassette was separated from the mold. The tissues were then sectioned using Leica ultramicrotome where the blade angle was set at 5^0^ and 10 µm sections were cut. The sections were floated on 37^0^C water bath on clean glass slides. The slides were kept at 65^0^C for 20 minutes for the tissue to adhere to the slide. These were then immediately stained.

**Staining:** The slides were then stained according to the outline given below-

- Xylene for 4 minutes
- Absolute ethanol for 4 minutes
- 95 % ethanol for 2 minutes
- Water for 2 minutes
- Hematoxylin for 3 minutes
- Water wash for 1 minute
- 95% ethanol for 1 minute
- Eosin 1 minute
- 95% ethanol for 1 minute
- Absolute alcohol for 2 minutes
- Coverslips with mounting media

The slides were then visualized under bright-field microscopy to check the epidermis, dermis, tissue integrity, number of hair follicles and sebaceous glands. The images were then analysed by ImageJ for determining the above parameters.

**Statistical Methods:** All the data analysis, mean, Standard Error of Mean (SEM) calculations and statistical significance tests were carried out using Origin Pro 9.1 Software. For statistical significance, Student’s t-test was used.

**Table S1: Hydrodynamic Diameter of Fullerene nanoformulation in DMEM+ 10% FBS with Concentration (700 rpm, 14 days).**

| **Concentration**  **(µg/mL)** | **Hydrodynamic Diameter (Z-av)**  **(nm)** | **Standard Error of Mean** |
| --- | --- | --- |
| Control | 0 | 0 |
| 0.01 | 15.625 | 1.20402 |
| 0.025 | 16.596 | 1.39209 |
| 0.05 | 17.30433 | 2.27788 |
| 0.075 | 18.193 | 2.05849 |
| 0.1 | 19.26 | 1.14136 |
| 0.5 | 19.73 | 1.6744 |
| 1 | 20.65333 | 2.46173 |
| 2.5 | 28.16 | 2.5349 |
| 5 | 34.37333 | 2.20621 |
| 7.5 | 43.31333 | 2.48201 |
| 10 | 47.26 | 3.298 |
| 25 | 82.5 | 9.278 |
| 50 | 132.1 | 9.618 |
| 100 | 153.7 | 12.345 |
| 200 | 175.4 | 11.687 |
| 300 | 196.62 | 12.167 |
| 400 | 221.1 | 21.42 |
| 600 | 353.8 | 26.15 |
| 800 | 554.1 | 36.047 |
| 1000 | 625 | 38.047 |
| 2000 | 765.3 | 35.623 |
| 3000 | 885.1 | 40.525 |
| 4000 | 890.4 | 27.829 |

**Figure S1**

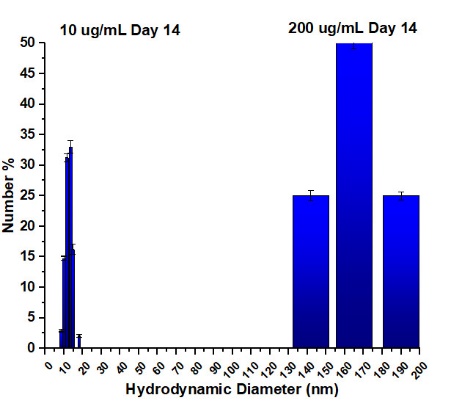

**Figure S1: The size distribution analysis (number % weighing) after 14 days for low concentration (10 µg/mL) and high concentration (200 µg/mL) showed predominant species of 15 nm and 170 nm, respectively.**

**Figure S2:**

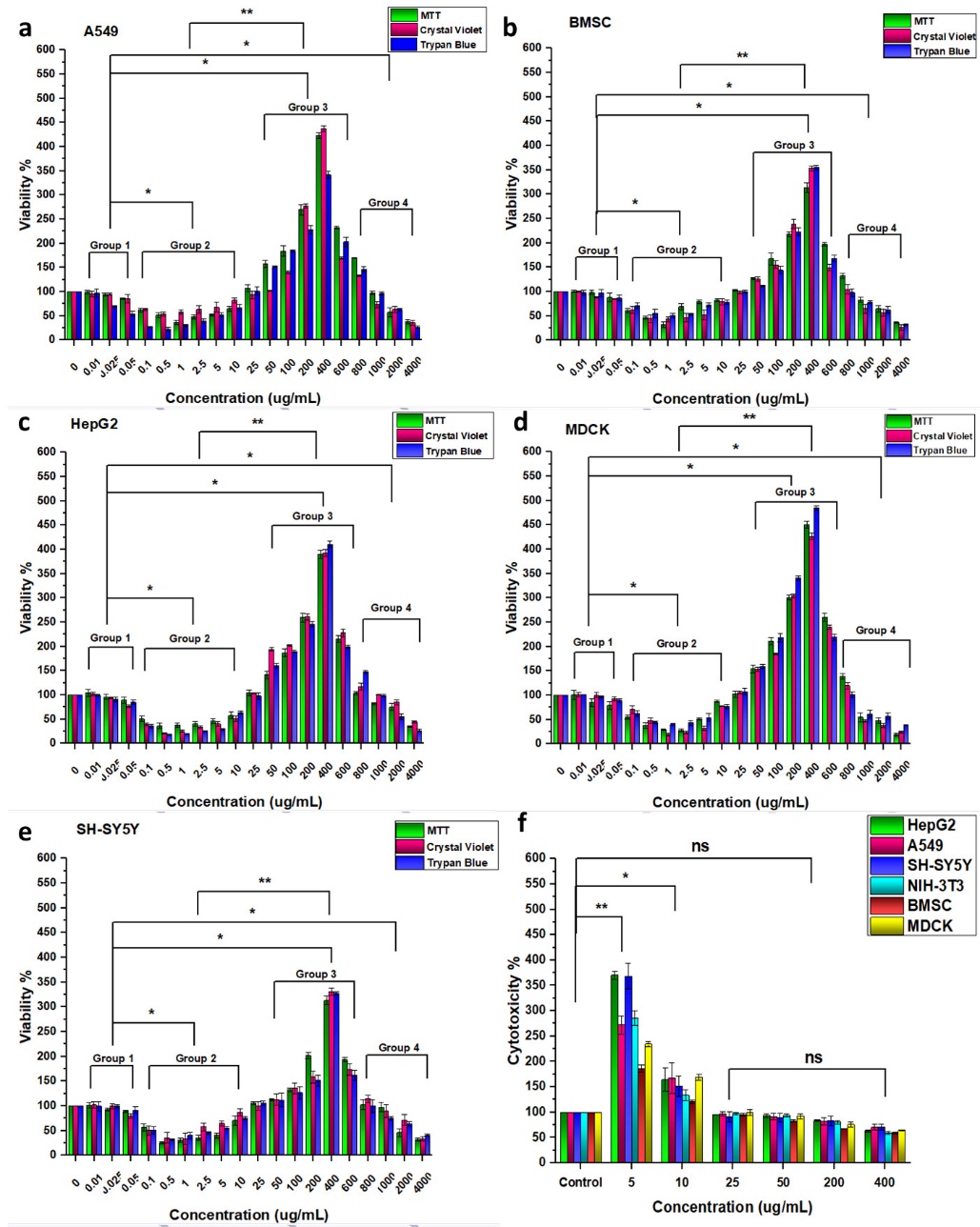

**Figure S2: (a-e) Cell viability trend on different cell lines with concentration of Fullerene. The cell viability is lower than control for 20-50 nm sized particles whereas viability increases for 130-230 nm sized particles. (Group1:** ~**20 nm), (Group 2:** ~**20-50 nm), (Group 3:** ~**80-230 nm), (Group 4:** ~**300-900 nm); (f) Cytotoxicity assay by LDH measurement. The concentrations showing lower cell viability (5 µg/mL (**~**35 nm) and 10 µg/mL (**~**50 nm)) in viability graphs (a-e) showed increased cytotoxicity in LDH assay in graph (f). On the other hand, the concentrations showing neutral or proliferative action on cells (25 µg/mL (~90 nm); 50 µg/mL (**~**120 nm); 200 µg/mL (~150 nm**) **and 400 µg/mL (**~**220 nm)) did not show any significant cytotoxicity effect in LDH assay in graph (f). (*p<0.05, **p<0.01)**

**Figure S3:**

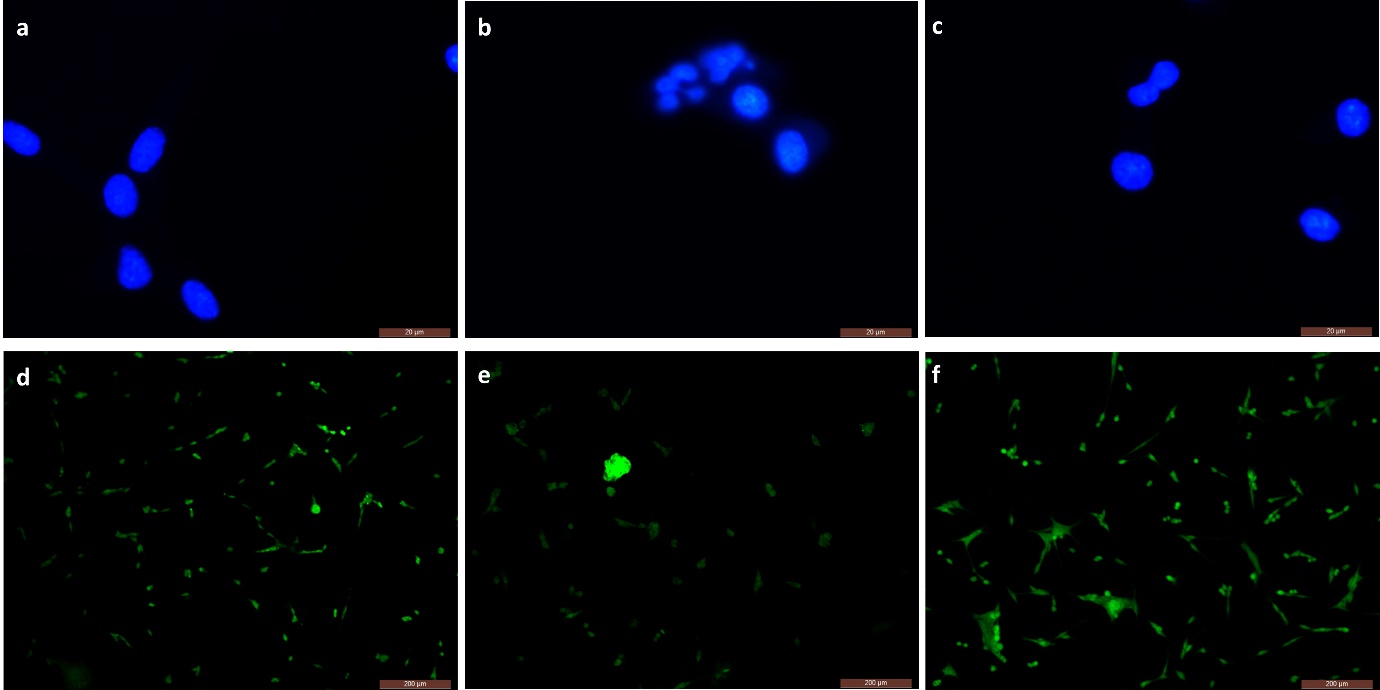

**Figure S3: DAPI and FDA staining of MDCK cells treated with (a, d) only media, (b, e) F_T30_, (c, f) F_P200_. F_T30_ treated cells show micronucleus, fragmented nuclei and lower number of viable cells. F_P200_ treated cells show mitotic bodies and a greater number of viable cells than control. (Upper panel scale bar 20 µm, Lower panel scale bar 200 µm)**

**Figure S4:**

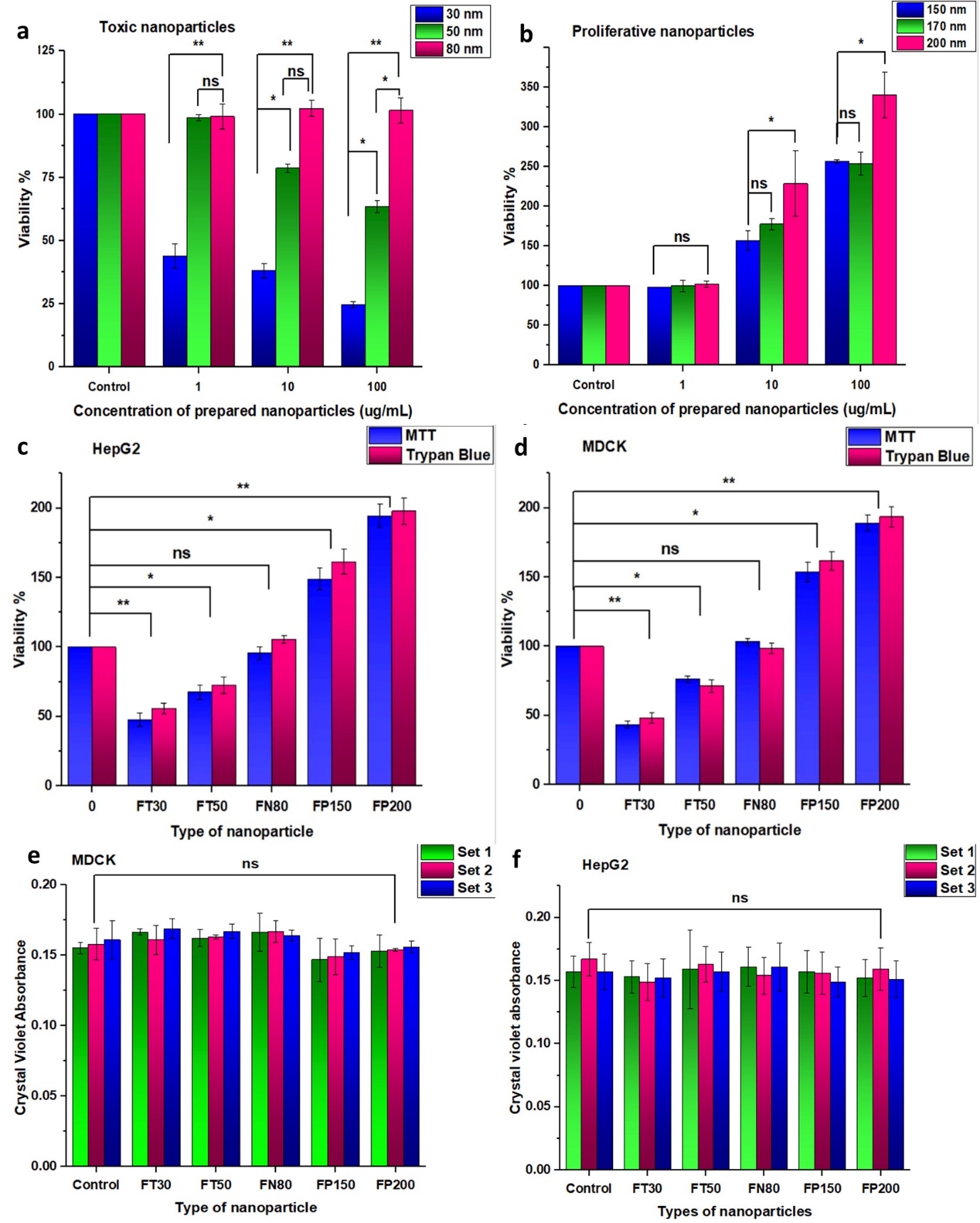

**Figure S4: (a,b) Deciphering size vs. concentration on MDCK. When prepared smaller nanoparticles of 30 and 50 nm are applied on cells, cytotoxicity was observed for all amounts of nanoparticles (1-100 µg/mL). However, for a particular size, the effect increases with the amount. For 80 nm sized particles, we observed similar viability as control in all the concentrations. Whereas when larger nanoparticles of 150, 170 and 200 nm are applied, increased viability was observed. The results suggest that size play the primary role in determining the biological activity. However, if we increase the amount of a particular size, the biological activity increases or decreases with the amount. (c,d) Effect of Fullerene nanoformulation (F_T30_, F_T50_, F_N80_, F_P150_, F_P200_) on the viability of 3D spheroids formed from (a) HepG2 and (b) MDCK; (e,f) The migration assay of the cell lines (a)MDCK and (b) HepG2 across 200 nm transwell filters. Set 1: The cells were treated with the nanoformulations (F_T30_, F_T50_, F_N80_, F_P150_ and F_P200_) after seeding on the filter to observe whether in response to the established concentration gradient, the cells migrate through the pores or not. Set 2: The cells were first treated with the respective nanoformulation treatment and then seeded on the filters. This set was performed to observe whether the treated cells migrate after coming in contact with normal media to check for transformed cells. Set 3: The lower chamber has media along with nanoformulation but the cells were not directly treated. This set was to check the effect of the nanoformulation as a chemotactic agent on the cells. (*p<0.05, **p<0.01, ***p<0.005)**

**Figure S5**

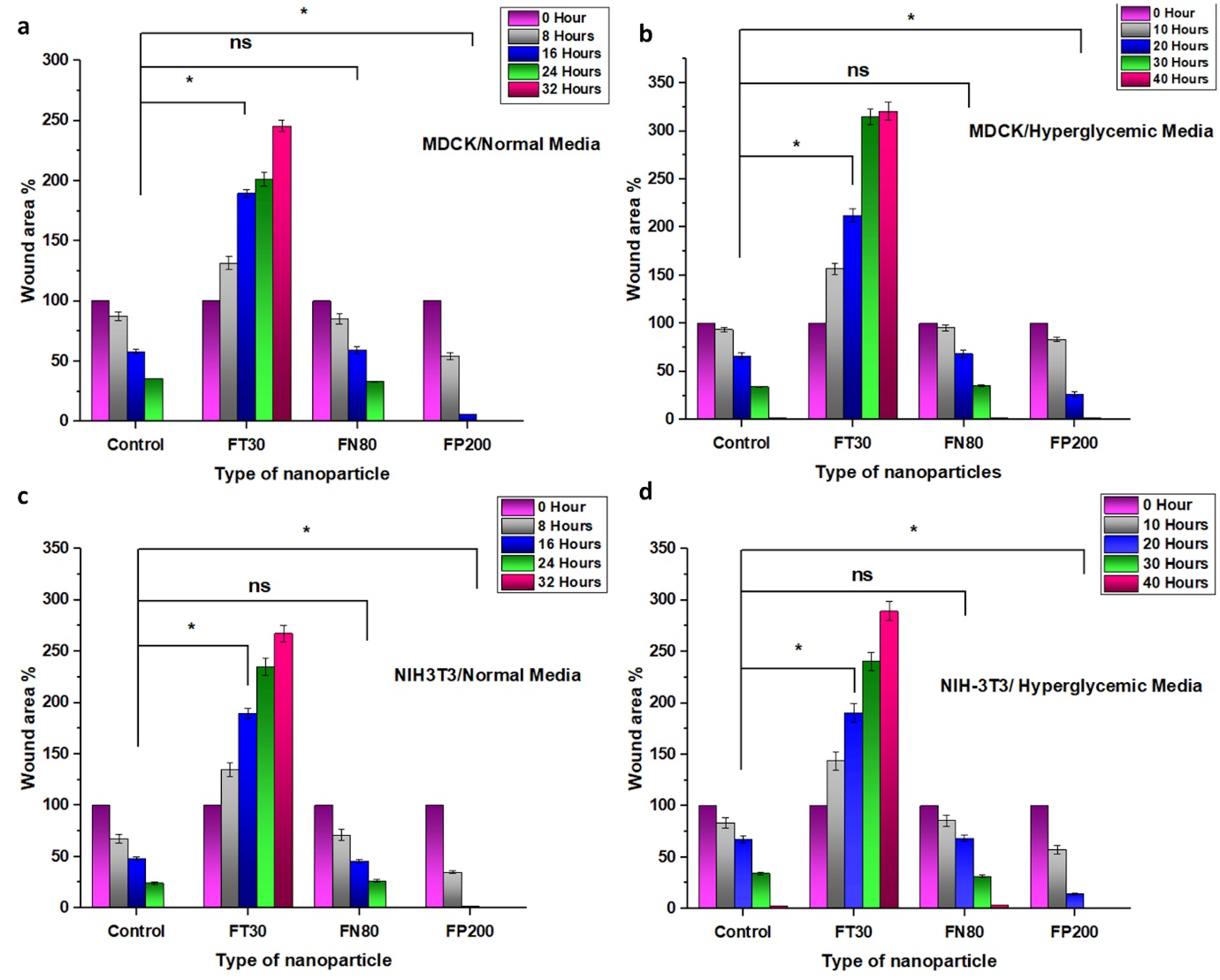

**Figure S5: Scratch assay quantification of cellular wounds after 3 types of nanoformulation treatment (Toxic, Neutral, Proliferative) by ImageJ. (a, b) MDCK scratch assay in normal and hyperglycemic media. (c, d) NIH-3T3 scratch assay in normal and hyperglycemic media. Normal media: The control and F_N80_ particles led to the wound closure in the same pace. In F_T30_, the wound area % increased with time in contrast to F_P200_ where wound area closed at an accelerated pace in both cell lines. Hyperglycemic media: The control and F_N80_ treated cells showed complete recovery in 40 hours compared to the F_P200_ accelerated effect where wound closed in 30 hours. The F_T30_ showed wound area expansion throughout 40 hours. (*p<0.05) (Number of wounds=9 for each treatment)**

**Figure S6**

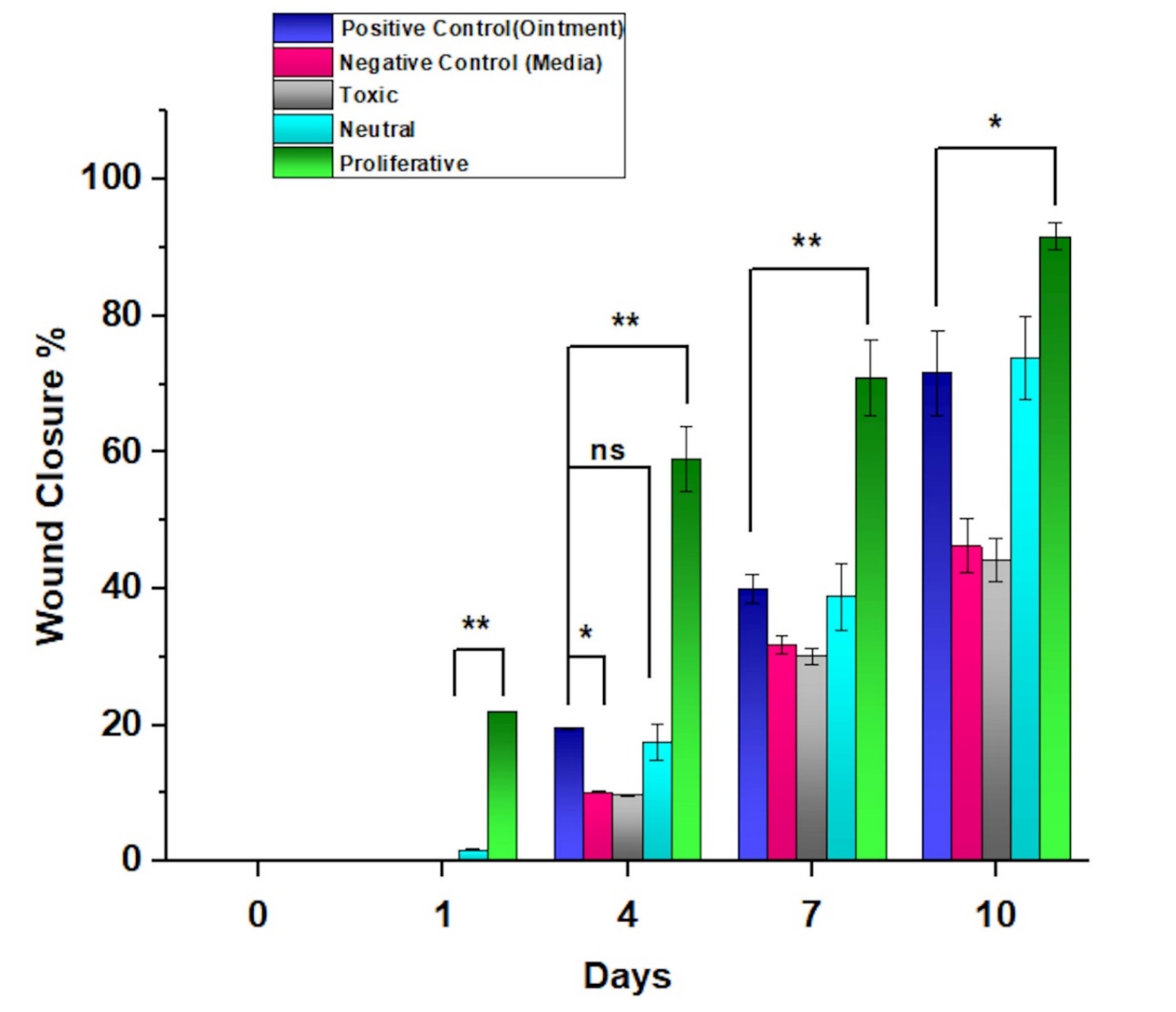

**Figure S6: *In-vivo* quantification of wound closure over 10 days. The Negative control and Toxic treatments show the slowest repair followed by Positive control and Ointment which shows intermediate repair ability. The Proliferative treatment shows the fastest repair (*p<0.05, **p<0.01). Number of wounds=4 for each type of treatment.**

**Figure S7:**

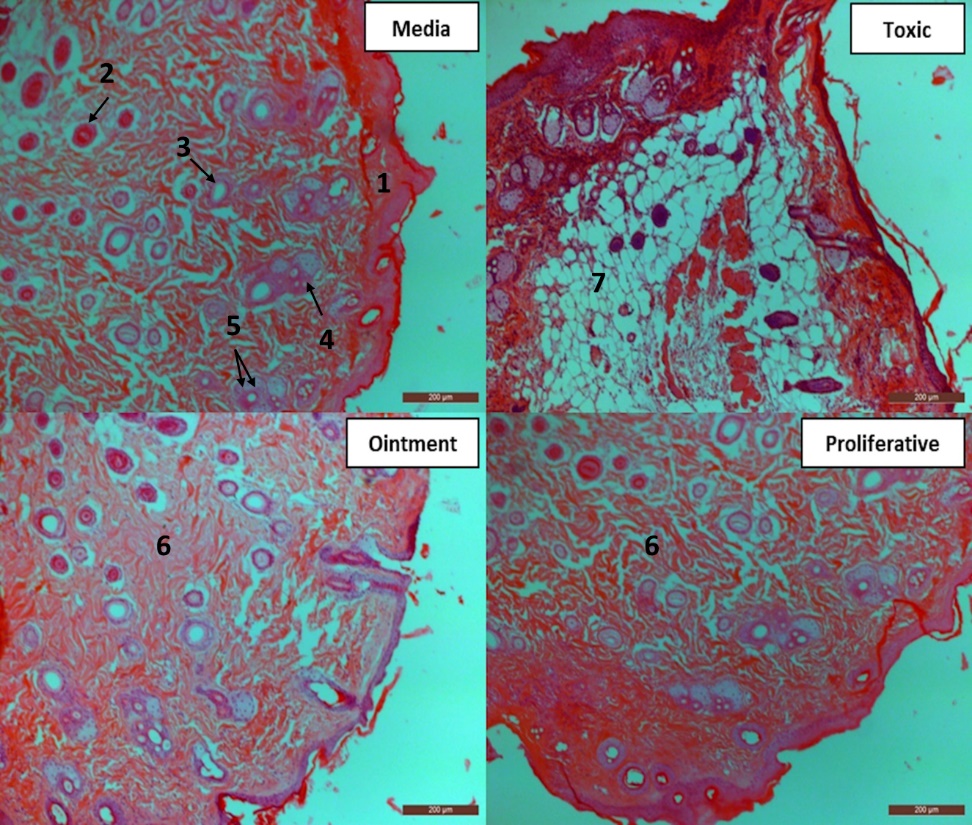

**Figure S7: Hematoxylin and eosin staining of the tissue sections of BALB/c after 14 days of complete healing. [1]: Epidermis with stacked appearance. The epidermis is of uniform thickness in all samples except Toxic treatment where it is significantly thicker. [2]: Blood capillary sections. All sections show blood vessel formation except the Toxic treatment. [3]: Hair follicles. The number of hair follicles are similar in Media and Ointment treatments, absent in Toxic and more numerous in Proliferative. [4]. Sebaceous glands. Uniform in number in all sections except Proliferative where these are more numerous. [5]. Inflammatory cells stained purple by Hematoxylin. These are uniform in number in all sections. [6]. Uniform matrix deposition observed in all sections except Toxic. [7]. Poor and discontinuous tissue matrix deposition in Toxic treatment. All elements are analysed and quantified by ImageJ. (Scale bar 200 µm)**
